# Land-use changes impact root-fungal network connectivity in a global biodiversity hotspot

**DOI:** 10.1101/2024.10.05.616733

**Authors:** Carina Carneiro de Melo Moura, Nathaly R. Guerrero-Ramirez, Valentyna Krashevska, Andrea Polle, Iskandar Z. Siregar, Johannes Ballauff, Ulfah J. Siregar, Francisco Encinas-Viso, Karen Bell, Paul Nevill, Oliver Gailing

## Abstract

1. Cross-kingdom associations play a fundamental role in ecological processes. Yet our understanding of plant-fungal co-occurrences in tropical rainforests and the potential impacts of land-use change shaping species connections remains limited.
2. By using amplicon sequencing on DNA from roots and their associated fungal communities, we aim to understand the impact of rainforest transformation on the composition and structure of root-fungal ecological networks in human-modified landscapes in Sumatra, Indonesia.
3. Each land-use type supports a distinctive set of indicator species, which are organisms that reflect specific environmental conditions and can signal changes in ecosystem health. We observed a decline in the richness of plant species indicators and plant-fungal associations with increasing land-use intensification. Additionally, there is a turnover in root communities, shifting from native and endemic species in rainforests to non-native, generalist herbaceous species in rubber and oil palm plantations.
4. Plant-fungal connectivity significantly declined with increasing land-use intensification, suggesting that managed ecosystems may have weakened root-fungal interactions. Network analysis highlights the distinct responses of various fungal groups. For instance, arbuscular mycorrhizal fungi (AMF) showed fewer connections with modules linked to oil palm and rubber roots, indicating weakened root-fungal associations in monocultures. This aligns with the observed reduction in AMF diversity in converted land-use areas compared to forests, further reinforcing the negative impact of land-use practices in oil palm and rubber monocultures on AMF diversity.
5. Synthesis. Dimensioning the impacts of rainforest transformations belowground is constrained by our understanding of fungal functional guilds. Highly modified systems exhibited fewer connections, suggesting a dynamic restructuring of root-fungal relationships in response to land-use changes. Understanding the intricate interplay between plants and fungi in the face of land-use change can provide valuable information for conservation efforts, agricultural practices, and ecosystem management strategies aimed at promoting biodiversity, soil health, and ecosystem resilience in the context of changing environmental conditions. Moreover, it underscores the importance of communities’ networks in land-use planning and management decisions to support plant and fungal diversity in terrestrial ecosystems.

## Introduction

Tropical rainforests are highly diverse ecosystems that contain a wide array of micro-habitats and organisms^1,2^. Yet, rainforests face major threats due to their rapid replacement with cash crop plantations such as oil palm and rubber^3,4^. For example, the ongoing agricultural expansion has resulted in ∼21 million hectares of oil palm plantations globally^5,6^. Overall, the taxonomic diversity of native species has decreased due to land-use changes^7–11^. However, to better understand the multiple impacts of agricultural expansion on tropical ecosystems, we urgently need to strengthen our knowledge of impacts of land-use conversion on species interactions^7–9,12^.

Plant-fungal associations play a fundamental role in shaping the structure and functioning of tropical ecosystems^6,13,14^, providing insights into the resilience, adaptive capacity, and health of forest ecosystems^15^. Root-microbial interactions involving different kingdoms and functional guilds^15–19^ have been linked to mutualistic preferences, evolutionary and trait differences among hosts, and phylogenetic relatedness and competitive exclusion between fungi^20,21^. For example, the abundance and diversity of arbuscular mycorrhizal fungi (AMF) have been found to respond to changes in plant diversity and composition^22,23^, with host-AMF preferences likely modulated by plant functional groups, instead of reflecting individual species interactions^21,24,25^. Moreover, an increase in the abundance of AMF in the rhizosphere is expected to promote a decrease in pathogen abundance^21,25,26^ and to be positively correlated with the abundance of saprotrophic fungi^21,27^. Although in tropical lowland forests, most tree species are associated with AMF, plant species in the Dipterocarpaceae and Fagaceae families are known to harbor ectomycorrhizal fungi (ECM)^28,29^. Nevertheless, our comprehension of root-fungal associations in lowland tropical rainforests stems predominantly from studies concentrating on root-AMF^23,30^, with EMF studies mainly from temperate ecosystems^31–34^.

Large-scale conversion of tropical rainforests to monocultures reduces microhabitat diversity, leading to species turnover^8,10,11^ and loss of plant and fungal diversity^10^. For example, when analyzing species-specific root-AMF associations in individual root samples, there was a significant decrease in AMF richness in roots of oil palm and rubber compared with roots from plants sampled in rainforests^24^. Environmental filters associated with forest conversion may select tolerant and opportunistic species (plants and fungal) that can occupy large niche ranges and outcompete native species in converted landscapes^17,35–39^, resulting in changes in community composition. Therefore, insights toward comprehending community-level interactions in natural and human-modified ecosystems are highly needed^40–42^, yet studies in tropical diversity hotspots on belowground association are rare.

By combining amplicon sequencing on DNA from identical samples for plant and fungal communities’ assessment with network analysis, we can provide valuable insights into the mechanisms driving the coexistence of species from various kingdoms within biological communities, revealing trophic and non-trophic interactions under specific environmental conditions^3,43–45^. Network analysis plays a crucial role in identifying highly connected nodes, facilitating the identification of key species and interactions essential for shaping community structure^13,27,46^. An effective approach for identifying land-use-specific species (i.e., indicator species), is the use of the indicator value index (IndVal), which evaluates the distribution patterns that best match the taxa in question^47^, determining whether a species is specific to a particular land-use or associated with multiple land-use types.

Here, by using an unprecedented dataset for the tropics containing roots and fungal communities from four land-use types in Sumatra, Indonesia, we aim to understand the impact of rainforest transformation on the composition and structure of root-fungal networks in a global tropical hotspot. To do that, we first identified patterns in root-fungal communities and estimated indicator species associated with each land-use type. Second, we estimated root-fungal associations through co-occurrence network analysis. Third, we examined highly connected taxa that shape the composition and structure of ecological networks. We hypothesized that forest transformation is a main driver of root-fungal associations via changes in root diversity, composition, and traits^11^. Plantations, for instance, have 59% fewer plant species than rainforests, where the latter consists predominantly of native species, including forest-specialist, versus higher non-native species richness and abundance in other land-use types^4^. Thereby, we expected rainforest plots to be associated with roots from rare and endemic species. In contrast, roots from non-native and generalist species, along with fungal groups adapted to withstand or thrive in highly disturbed niches, may serve as indicators of land-use intensity due to their resilience to adverse conditions. Finally, across land-use types, we hypothesized that taxa associated with rainforests have greater ecological connectivity than those in plantations, thereby contributing to a more structured ecosystem.

## Materials and Methods

### Study site and sampling

Our study was conducted in the lowlands of Jambi Province, Sumatra, Indonesia, in the framework of the EFForTS project (Ecological and Socioeconomic Functions of Tropical Lowland Rainforest Transformation Systems). Two representative landscapes undergoing extensive changes in land use and containing remnant forest areas were selected, Bukit Duabelas National Park and Harapan Rainforest (Fig 1). The region has a tropical humid climate with two distinct seasons: a rainy period spanning from December to March, and a dry period typically occurring around July and August. The average annual temperature is 26.7 °C, and the region receives an average annual precipitation of 2235 mm. The area has experienced significant rainforest conversion into agricultural systems, primarily for cultivating cash crops such as oil palm (*Elaeis guineensis*) and rubber (*Hevea brasiliensis*)^4^. The study area is predominantly characterized by natural vegetation, specifically dominated by trees belonging to the family Dipterocarpaceae^48^.

**Figure 1.**
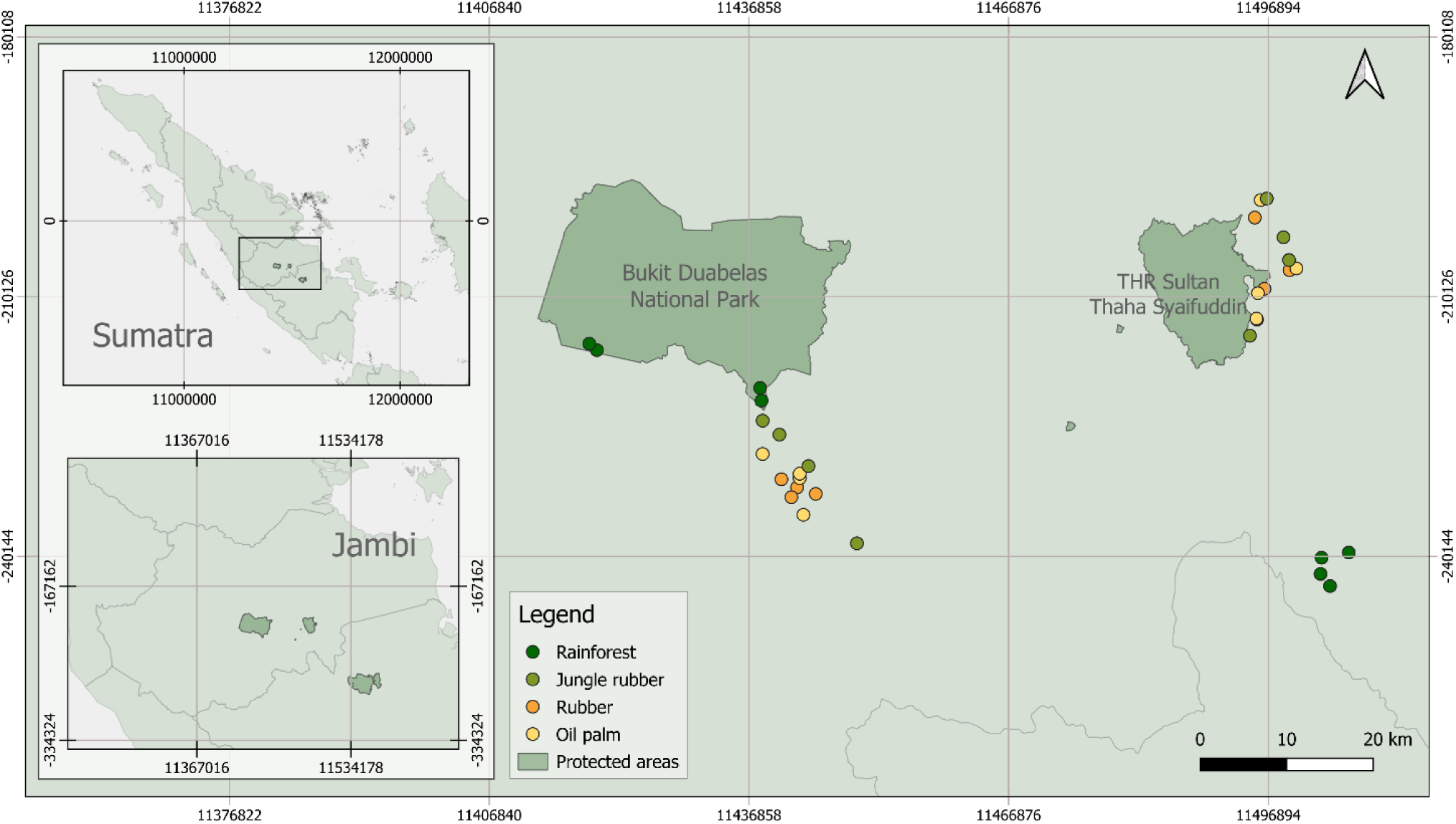
Plots are located in two landscapes, Bukit Bukit Duabelas and Harapan Rainforest (which includes THR Sultan Thaha Syalfuddin), at the Jambi district in Sumatra, Indonesia. Plots represent four land-use types in the region, i.e. rainforest, jungle rubber, rubber, and oil palm plantations.

Within the EFForTS project, 32 plots of 50×50 m across four land-use types were established, with eight plots per land-use type, namely logged-over old-growth rainforest, jungle rubber agroforestry, and rubber and oil palm monocultures. In each randomly selected 5×5 m subplot installed within each plot, five soil cores with a diameter of 0.04 m and 0.20 m depth were obtained from the four corners of the subplots, as well as one at the center in 2016. The samples were placed in a cooling box during the field sampling and then stored at 4° C at the University of Jambi. The five cores per subplot were pooled, sieved, and sorted as fine or coarse roots using two sieves with 5-and 10-mm mesh and freeze-dried prior to DNA isolation of root samples as described in Ballauff et al.^11^.

### Molecular and bioinformatic procedures

In this study, we used a DNA metabarcoding approach on the selected root samples to identify the plant and fungal species. However, two samples—one from oil palm and one from rubber plots—were excluded from the analysis due to insufficient quantity (representing n = 30 plots). Dried fine roots were shredded in a RETSCH Mixer Mill MM 400 at 30 Hz for 20-30 seconds until powdered, kept in 2 ml tubes and maintained in liquid nitrogen during processing. DNA isolation of fine roots was done using the innuPREP Plant DNA Extraction Kit (Innuscreen GmbH) by re-suspending the samples in 100 µl of water and using 400 µl of the Lysis Solution SLS following the recommendations of the manufacturer (Innuscreen GmbH) and replacing the elution buffer with nuclease free water for the last step.

Templates of root DNA were amplified using the ribulose-1,5-bisphosphate carboxylase large chain (*rbcL*) region with the primers rbcL2 5’-TGGCAGCATTYCGAGTAACTC-3’^49^ and rbcLa-R 5’-GTAAAATCAAGTCCACCRCG-3’^50^, which yields PCR products of approximately 450 bp in length. Triplicate PCRs were conducted to enable the detection of plant species with low DNA concentrations. Each reaction contained a final volume of 14 µL using 6.8 µL H_2_O, 1µL DNA (10-20 ng/µL), 1.5 µL of 10X PCR Buffer (with 0.8 M Tris-HCl, 0.2 M (NH_4_)_2_SO_4_), 1.5 µL of MgCl_2_ (25 mM), 1 µL dNTPs (2.5 mM of each dNTP), 1 µL of each forward and reverse primers (5 pmol/µL) and 0.2 µL of Taq Hot FirePol (5 U/µL) from Solis BioDyne (Estonia). Positive and negative controls were executed alongside each PCR run, and no product detection was observed in the negative control samples. The thermal cyclic settings were programmed to an initial denaturation step at 95 °C for 15 min, followed by 35 cycles at 94 °C for 1 min, 50 °C for 1 min, 72 °C for 1 min, and a final extension step of 72 °C for 20 min. The PCR products were purified using the GENECLEAN Kit (MP Biomedicals) and pooled before library preparation. Concentrations were measured using a Qubit fluorescence spectrophotometer (Life Technologies) and standardized to 200 ng per sample. The Illumina TruSeq Nano DNA High Throughput Library Prep Kit (96 samples), and Illumina TruSeq DNA CD Indexes were used to prepare sequencing libraries. Paired-end sequencing was carried out on an Illumina MiSeq using the MiSeq Reagent kit v2-300 cycles with DNA sequencing libraries at 10 pM with 10% PhiX control at NIG (NGS Integrative Genomics Core Unit at the University of Göttingen).

The fungal community data identified using the ITS region were obtained from Ballauff et al.^11^, representing a subset of the dataset from the same study sites. The primer regions ITS1-F_KYO2 5’-TTYRCTRCGTTCTTCATC-3’^51^ and ITS2 5’-GCTGCGTTCTTCATCGATGC-3’^52^ including overhanging Illumina adaptors were used for amplification of the ITS 1 and 2 regions targeting the fungal community. For each DNA extraction obtained from fine root samples, PCR was executed using 0.25 μl of Phusion High-Fidelity DNA Polymerase (2 U μl ^-1^), 5 μl of 5x Phusion GC buffer (Thermo Fisher Scientific, Waltham, USA), 0.075 μl of MgCl_2_ (50 mM), 1.25 µl of DMSO (5%), 1.25 µl of bovine serum albumin (8 mg ml ^-1^), 0.5 μl of dNTP mix (10 mM each, Thermo Fisher Scientific, Osterode am Harz, Germany), 0.5 μl of each primer (10 mmol/l, Microsynth, Wolfurt, Austria) and 2 μl of template DNA (5 ng μl ^-1^). The PCR mix was adjusted to a total volume of 25 μl with dNTP free water. The cycling parameters were 1 cycle of 98 °C for 30 s, 30 cycles of 98 °C for 10 s, 47 °C for 20 s, and 72 °C for 20 s, and a final extension at 72 °C for 5 min. Three technical replicate reactions were performed and pooled before sequencing.

Quality control of the Illumina raw reads obtained in this study was done using FastQC^53^. Sequences were trimmed to remove primers and adapter sequences using Cutadapt^54^. We used Usearch 11.0.667^55^ to merge forward and reverse reads using the command -fastq_mergepairs. Low quality sequences (<Q score 20, <100 bp, ambiguous base-pairs) and singletons were removed using the commands -fastq_filter and -minsize. Sequence reads were de-replicated (-fastx_uniques), sorted by size (-sortbysize), and clustered (-cluster_otus) using the algorithm UPARSE-OTU^56^.

We opted to employ Operational Taxonomic Units (OTUs) to mitigate noise and minimize the impact of sequencing errors^56^. All OTU sequences were then compared against the NCBI Genbank database to verify the accuracy of the OTU assignments using a Blastn search, assuming the lowest common ancestor that has the highest sequence similarity value as the correct assignment^48^ and using 97% as similarity level, which allows assignment at least to the genus level. Based on the existing literature, plant root OTUs were categorized either as native or non-native. Additionally, they were classified based on their life form, such as tree, shrub or herb^4,57^ (http://www.plantsoftheworldonline.org/).

Fungal OTUs were categorized based on the trophic modes as saprotrophic, pathotrophic and symbiotrophic fungi by employing the FUNGuild annotation tool (https://github.com/UMNFuN/FUNGuild)^58^, while non-assigned sequences were labelled as unidentified.

A total of 1,916,098 paired-end reads were obtained using the *rbcL* assay for roots, ranging from 35,948 to 91,265 sequences per sample. The ITS region targeting fungi retrieved a total of 1,013,513 paired-end reads, ranging from 5,762 to 76,331 reads per sample.

### Downstream analysis

All data visualization and data analysis were conducted using rarefied OTU sequence counts for roots and fungi separately based on rarefaction depth using minimum sample depth in the package Phyloseq^59^ in RStudio, R version 4.2.2^60^. The dissimilarity among samples was calculated using a weighted UPGMA tree based on UniFrac distances between samples with the function Unifrac in the R package phyloseq^59^. We tested for differences in root and fungal communities detected in the four land-use types using a Permutational Multivariate Analysis of Variance (PERMANOVA) calculated by pairwise comparisons of the Unifrac distances in the R package vegan^61^. To assess the presence of spatial autocorrelation in fungal and root diversity within each land-use type, we employed Moran’s I statistic. Specifically, Moran’s I statistic evaluates whether the spatial distribution of diversity is more clustered than expected by chance. This analysis was performed using the R package spdep^62^. Moran’s I was calculated for different land-use types by creating spatial weights matrices based on nearest neighbors, and then testing for significant spatial clustering in the diversity data.

To pinpoint links between land-use and species composition, we identified root and fungal taxa associated with single or multiple land-use types. This was achieved by implementing the IndVal.g function from the R package indicspecies^63^ using 10^4^ permutations. OTUs were assigned as indicators for specific land-use systems based on point-biserial correlation coefficient (r), considering results significant at p < 0.05. This approach estimated the correlation coefficient (r) of an OTU’s positive association with one or more land-use types. The IndVal index calculates the highest association values between species and each site group or combination of site groups^47^. To visualize composition of significant (p < 0.05) OTU associations (indicator taxa) to each land-use type, bipartite networks for each taxonomic group (plants and fungi) were employed using the Fruchterman-Reingold layout with 10^4^ permutations in the R package igraph^64^. Furthermore, we tested for differential OTU abundance across land-uses using likelihood ratio tests (LRT) in the package edgeR^65^, following Hartman et al.^66^.

We constructed and visualized associations between root and fungal OTUs and the land-use systems focusing on plant and fungal indicator species. To do that, the plant-fungal network was calculated using significant correlations, p-value < 0.001 and Spearman correlation coefficients (ρ) > 0.7, between root OTUs and fungal OTUs using 9999 permutations and plotted using the Fruchterman-Reingold layout algorithm in igraph^64^. Network metrics including the number of nodes, edges, and mean node connectivity were obtained using igraph^64^.

We characterized the distribution of OTUs in the network and their interactions within and between modules using the modularity analysis by simulated annealing^46^ employed in the R package rnetcarto^67^ using the OTUs that showed significant Spearman correlations (p < 0.001, ρ > 0.7). Node roles within the network were determined through a comprehensive analysis of the relationship between the participation coefficient (*P*), reflecting the diversity of connections across the network’s communities, and the within-module degree (z), measuring the connectivity of each node within its module. This assessment, conducted using rnetcarto, resulted in the categorization of nodes into distinct roles based on their connectivity and participation levels. Specifically, nodes were classified as kinless when their edges were evenly distributed among all modules (*P* > 0.80), connectors if they exhibited numerous edges to other modules (0.62 < *P* ≤ 0.80), peripherals if they displayed the majority of their links within their module (0.05 < *P* ≤ 0.62), and ultra-peripherals if they presented all their edges within their module (*P* ≤ 0.05)^46^. This classification provides a nuanced understanding of each node’s role in the network, shedding light on its connectivity patterns and contributions to the overall structure of the network^46^.

## Results

### Sequence data and community structure overview

Among the identified root OTUs, 247 were classified into 25 orders, 47 families, 111 genera, and 124 plant species. These assignments comprised 13 OTUs categorized as herbaceous species, 56 as shrubs, and 178 as tree species. Furthermore, the sequence assignments revealed the presence of eight species that are non-native to Indonesia. In addition, our dataset identified 40 OTUs at the genus level, potentially indicating either an incomplete availability of sequences for Indonesian species in sequence databases or a lack of diagnostic differences at the species level with the *rbcL* barcode. The weighted UPGMA tree revealed discernible land-use effects on the community composition of roots (p < 0.005 for all, except between oil palm and rubber) (Fig S1 A).

For the fungal sequences, 14,049 OTUs were categorized into 177 orders, 430 families, and 1,159 genera. Within the assigned OTUs, 2,010 OTUs were classified as symbiotrophs, 2,362 as saprotrophs, 540 as pathotrophs, and 1,067 as part of multiple functional guilds. 8,070 fungal OTUs could not be assigned to any functional group. The weighted UPGMA tree for the fungal dataset indicates minimal effects of land-use change on fungal community profiles (*p* = 0.709) (Fig S1 B).

The Moran’s I spatial autocorrelation test using root Shannon diversity across different land-use types revealed a non-random distribution in rainforests, with Moran’s I statistic of 0.1547 (sd = 2.9804, p = 0.00144), indicating significant spatial clustering of diversity (p < 0.01). Conversely, spatial patterns in converted land-uses were consistent with random distribution (jungle rubber: Moran’s I=-0.218, sd = -0.9823, p = 0.837; oil palm: Moran’s I = 0.1941, sd= -0.4624, p = 0.6781; and rubber: -0.1989, sd = -0.5583, p = 0.7117).

Similarly, the spatial autocorrelation of fungal diversity across different land-use types indicated a marginally significant tendency towards spatial clustering in rainforest, with Moran’s I statistic of - 0.0075 (sd = 1.3545, p = 0.08778). In contrast, spatial patterns in other land-use types were consistent with random distribution (jungle rubber: Moran’s I = -0.1792, p = 0.6742; oil palm: Moran’s I= - 0.1985, p = 0.6837; and rubber: Moran’s I = -0.2239, p = 0.8425).

**Figure S1.**
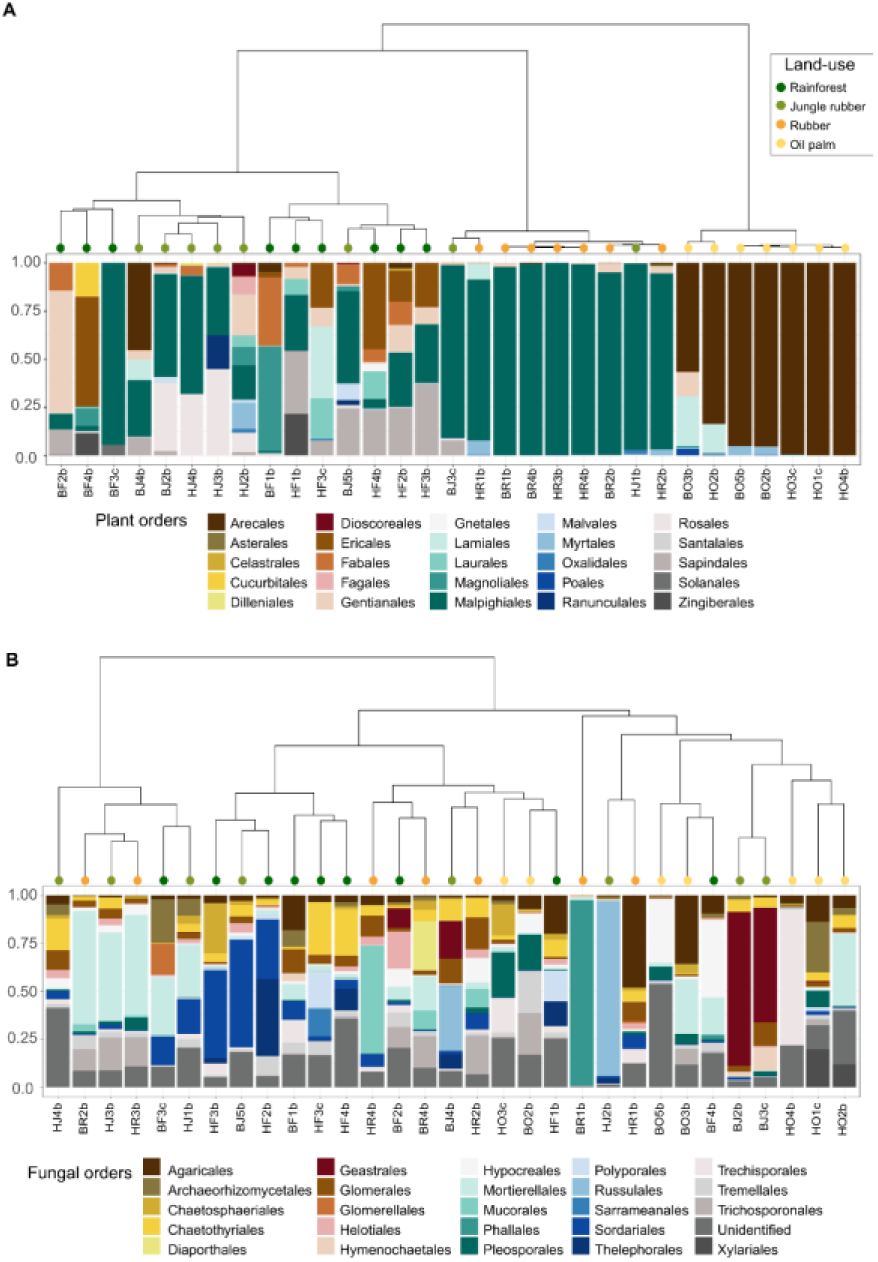
Plant and fungal community compositional similarity. Weighted UPGMA tree of all samples based on Unifrac distances and barplot of relative frequency of taxa composition for (A) plant OTU assignments, and (B) for fungal communities displayed by order. The colors of the tip labels depict land-uses (rainforest, jungle rubber, rubber, and oil palm monocultures). Labels depict names given to sample sites: H - Harapan Rainforest, B - Bukit Duabelas National Park, F - Rainforest, J - Jungle rubber, R - Rubber plantation, O - Oil palm plantation.

### Bipartite network of indicator species linked to land-uses

The diversity of plant indicator species and functional groups exhibited variations across land-use types (Fig 2, Appendix S1). Rainforests demonstrated a higher diversity of indicator species, encompassing a larger array of native and endemic taxa, as well as diverse plant life forms (Fig 2 A, B). In contrast, more intensive monoculture systems revealed the presence of non-native plant species such as *Asystasia gangetica* and *Clidemia hirta,* along with an increased number of other herbal species (Fig 2 C). Compositional distinctions were evident among land-use types for both root and fungal communities, with a higher similarity observed between less intensified land-use systems, such as rainforest and jungle rubber (Fig 2 A). Conversely, more intensified land-use systems such as monocultures featured fewer species identified as indicators. Indicator species of oil palm plantation plots included *Elaeis guineensis* (oil palm tree), alongside other indicator species such as the herbaceous species *Centotheca lappacea,* the non-native creeping herbs *Asystasia gangetica* and *Clidemia hirta*. In rubber monocultures, indicator species comprised *Hevea brasiliensis* (rubber tree), the native shrub climber *Ichnocarpus frutescens*, *Trachelospermum asiaticum*, and the herbs *C. lappacea*, and *C. hirta*.

**Figure 2.**
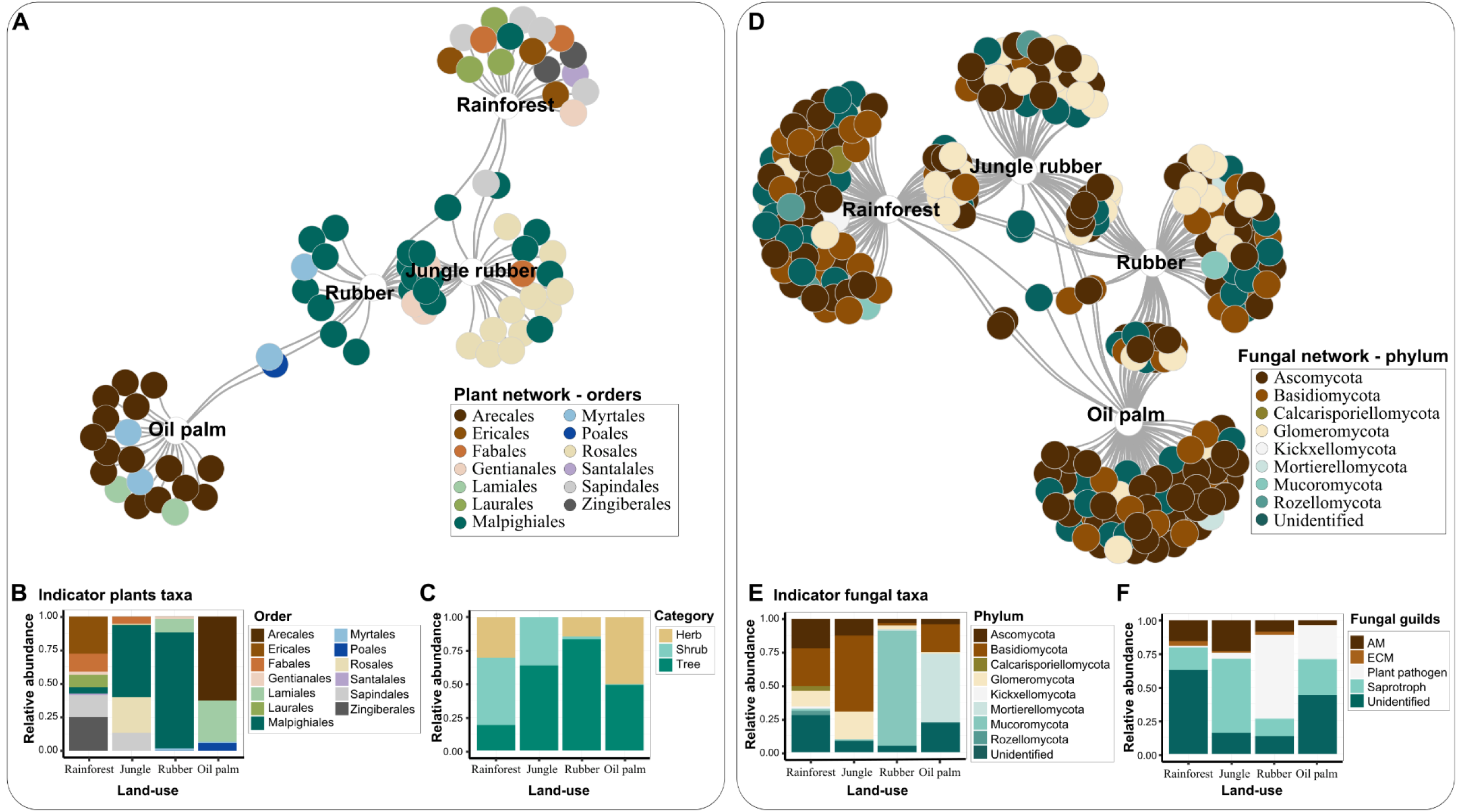
Bipartite networks of indicator species associated to the four land-use types: rainforest, jungle rubber, rubber, and oil palm monocultures. (A) Plant indicator species network at the order level. (B) Relative abundance of indicator plant roots in the four land-use types. (C) Relative abundance of life forms of indicator plant taxa in the four land-use types. (D) Bipartite network of fungal indicator species identified in root samples at the phylum level. (E) Relative abundance of indicator fungal taxa in the four land-use types. (F) Relative abundance of functional guilds of indicator fungal taxa in the four land-use types. The colors used in the figures correspond to the respective assignments of Operational Taxonomic Units (OTUs).

Land-use changes influenced the taxonomic and functional groups associated with fungal indicator species, as the composition and proportion of indicator species varied along land-uses (Fig 2). The fungal network illustrated a close association between the rainforest and jungle rubber clusters, while monocultures showed greater proximity to each other (Fig 2 D). The rainforest exhibited distinct fungal indicators from the phyla Calcarisporiellomycota and Kickxellomycota. In contrast, OTUs assigned to Mucoromycota and Mortierellomycota were prevalent as indicator groups in the monocultures (Fig 2 E). A substantial proportion of the reads associated with the monocultures among the fungal indicator taxa belonged to potential species-specific plant pathogens, while AMF identified as indicator OTUs were more abundant in less disturbed environments (Fig 2 F).

### Co-occurrence network between roots and fungi

The network analysis revealed that both modularity, i.e., partitions or clusters with denser connections, and connectivity, i.e., the degree to which nodes are interconnected, were influenced by land-use types and plant functional groups. The plant-fungal network consisted of 2,296 nodes (163 root OTUs and 2,133 fungal OTUs), and 47,922 edges (42,217 fungi-fungi, 580 plant-plant, and 5,125 plant-fungi) (Fig 3 A, Appendix S2). High modularity (Q = 0.9) indicates a strong community structure within the network, with simulated annealing optimizing the partitioning into 22 modules. The highest connectivity was observed in the module, which includes indicator species associated with rainforests with 16,418 edges, i.e., associations. In contrast, indicator species of jungle rubber had 2,981 edges, oil palm 2,883 edges, and rubber 203 edges. There were fewer significant plant-fungal associations when considering root OTUs assigned to herbaceous plant species (194), in contrast to the larger number of plant-fungal associations observed for trees (3,617) or shrubs (1,025) (Data S1). The disconnected nodes observed in the network (i.e., *Kunstleria ridleyi, Poikilospermum suaveolens* and *Suregada glomerulata*) may suggest that these species are either highly specialized, lack direct ecological interactions, or that some interactions were not detected due to incomplete data (Fig 3 A).

**Figure 3.**
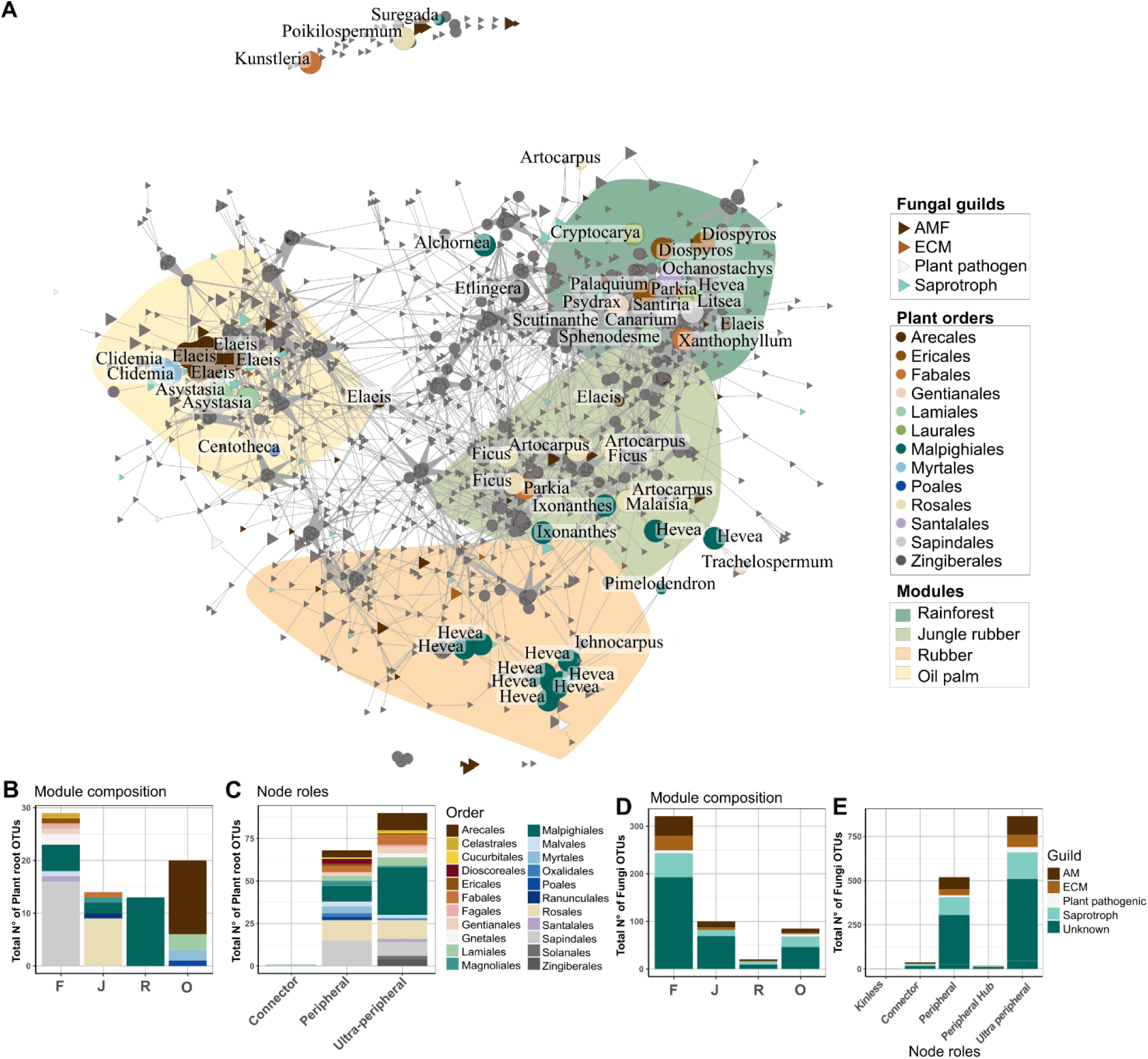
(A) Network based on significant co-occurrence correlations between plant roots and fungal. The displayed modules highlight clusters containing indicator plant OTUs linked to the four land-use types (rainforest, jungle rubber, rubber, and oil palm plantation), using the modularity analysis by simulated annealing. Circles show indicator plant OTU assignments presented at the genus level; colors depict the order of plant indicator OTUs. Triangles depict fungal indicator groups colored by functional guilds (AMF, ECM, plant pathogen, and saprotroph). (B) Plant composition for modules that included indicator plant OTUs, and (C) the taxonomic profile of node roles identified through the assignment of plant OTUs clustered in interkingdom modules. (D) Fungal composition for highlighted modules containing indicator fungal OTUs. (E) Taxonomic profile of node roles identified through the assignment of fungal guilds clustered in interkingdom modules. Colors correspond to the assignment of plant root OTUs at the order level and fungi at the guild level. N° =Number.

### Network topological roles

The network primarily comprised nodes categorized as peripheral, with 68 plant OTUs and 712 fungal OTUs, or ultra-peripheral, with 94 plant OTUs and 1,327 fungal OTUs (Fig 3 A). In addition, 73 nodes (one root and 72 fungal OTUs) were identified as connectors (Fig 3 and Fig 4, Appendix S1). Figures 3 B-E illustrate the taxonomic composition of modules associated with indicator species linked to the four land-use types. Among these modules, indicator species of rainforest exhibited the highest diversity and OTU abundance (Fig 3 B). Additionally, these rainforest-associated modules displayed the greatest proportion of AMF, ECM, and Saprotrophic fungi (Fig 3D). Species occupying peripheral and ultra-peripheral roles encompassed various taxonomic groups or functional roles, as illustrated in Fig 3 C and Fig 3 E.

**Figure 4.**
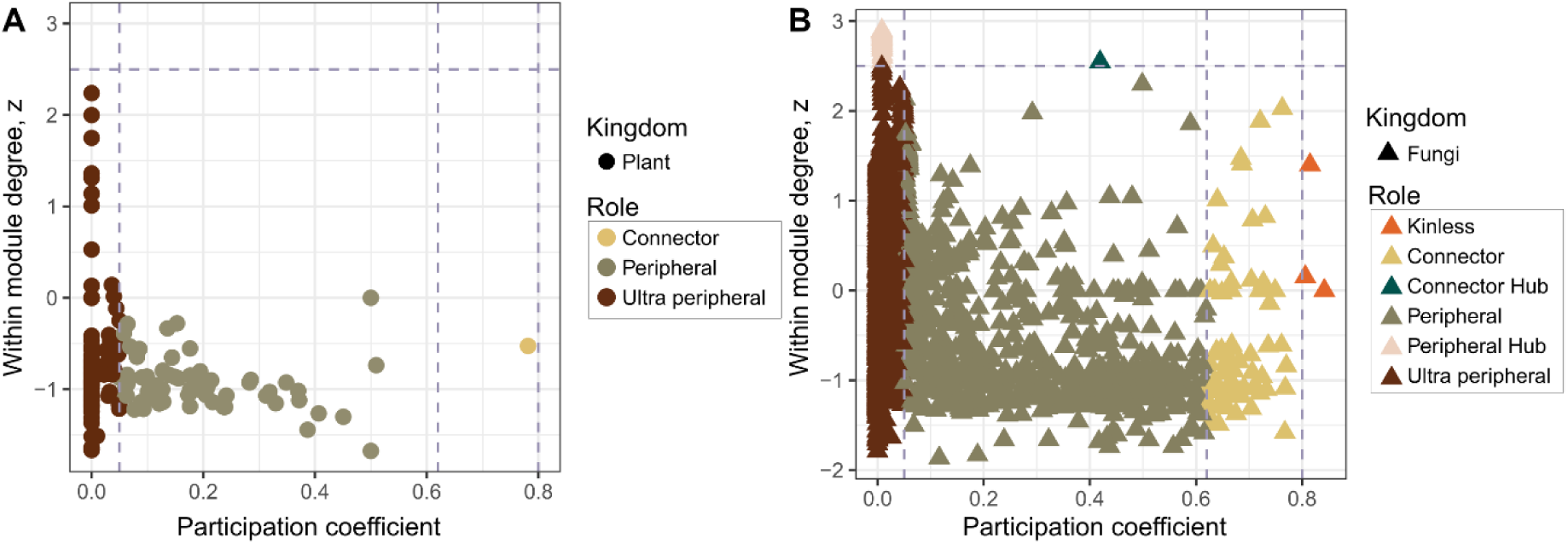
Topological node roles based on the distribution of the module degree (z) and participation coefficient (*P*) of each OTU. (A) Circular symbols represent plant OTUs, and (B) triangular symbols represent fungal OTUs. Colors correspond to the node role. The threshold of each node category is represented by traced lines. The within-module degree z[i] quantifies the extent of connectivity of each node [i] within the same module. Higher z[i] values indicate stronger within-module connections. The participation coefficient gauges the distribution of node’s [i] links across various modules; it approaches 1 if links are uniformly distributed across all modules and 0 if all links remain confined within their own module. Nodes with a z-score ≥ 2.5 are categorized as module hubs, whereas nodes with a z-score < 2.5 are classified as non-hubs.

The fungi *Dendrosporium* sp. and *Gerhardtia highlandensis* were classified as connectors and indicators of the rainforest system. Glomeraceae sp. and Herpotrichiellaceae sp. were indicators of the jungle rubber system and also played a role as connectors. These taxa are associated with environments presenting different degrees of disturbance. In oil palm and rubber, connectors and indicators included detritivores or taxa associated with oligotrophic conditions, such as Ascomycota sp., *Metschnikowia* spp., Pezizales sp., *Setophoma terrestris*, Herpotrichiellaceae sp., *Conlarium* sp., Sordariomycetes sp., *Ochroconis* spp., *Cladophialophora* spp., and Pezizomycotina sp. Glomeromycota sp.

Within the network, only three nodes were classified as kinless due to their high connectivity. Two nodes identified as ECM, *Pachyphlodes melanoxantha* and *Elaphomyces favosus*. The third node belonged to the category of unknown fungi. It is noteworthy that the diversity of indicator fungi occupying peripheral or ultra-peripheral roles was extensive and encompassed a range of functional groups (Appendix S2 and Data S2).

## Discussion

### Patterns in plant-fungal communities

Our findings align with the notion that the assembly of root fungal communities is influenced by land-use change^11,68^. For example, previous studies have demonstrated that rainforest conversion resulted in substantial shifts in the composition of soil fungal communities, despite a significant decrease in diversity not being observed^8,11^. The impact of intensive land-use practices and the consequent modification of abiotic and biotic filters in microhabitats are legacy effects of preceding land-uses on the composition of microbiome communities^12,68,69^. A reduction in root diversity can lead to a reduction in the availability of resources and ecosystem functions, as multiple species may rely on specific types of roots for their growth and survival^13,70^. Further, the disruption of ecological interactions such as plant-fungal associations influences nutrient uptake and plant growth, which may, in turn, impact herbivores, pollinators, and other species^71–73^.

The difference in the spatial distribution of plant and fungal diversity between rainforest and converted land uses reflects that in rainforests, rare and endemic species are more clustered spatially than randomly distributed. Non-random distribution of diversity suggests a healthy ecosystem, rich in habitat heterogeneity and species interactions^74^. In contrast, homogeneous habitats, as observed in converted land uses, indicate habitat simplification and a loss of niche diversity and ecological processes. Considering non-random distribution in rainforests is pivotal to understanding not only the impacts of forest conversion but also conservation and restoration^44^.

The 111 plant genera detected in this study using DNA metabarcoding account for approximately 20% percent of the genera previously documented in the same area using morphology-based plant inventories conducted by our research consortium over an 18-month sampling period^4^. It is worth noting that studies focusing solely on individual specimens of fine roots conducted in tropical forests or biodiversity experiments face challenges in capturing the same number of plant OTUs richness as observed in our study^21,23,24,30^. Therefore, a more complete coverage of root and interaction diversity using DNA metabarcoding in hyper-diverse regions under current global change pressures might benefit by increasing the number of soil samples. Although 20% may seem low in terms of capturing the full diversity of genera, it is important to recognize that the method used in our study provides a rapid snapshot of the landscape profile. This can be valuable for quickly assessing general trends in plant diversity. We acknowledge, however, that increasing the number of soil cores or sampling events would likely yield a more comprehensive representation of the species composition in the area. This approach serves as an initial overview, with the potential for more detailed studies to follow.

### Root-fungal network structure

The high degree of network compartmentalization observed in our network analysis indicates specific connections between community assembly of roots and fungi^13,75^. A decrease of host species in monocultures is often associated with the occurrence of stress-tolerant and opportunistic species^4,36,38^, which in our study was represented by opportunistic species of both the plant (e.g., *C. hirta* and *A. gangetica)* and fungi (*Cladophialophora* sp.). These generalist species thrive under low-resource conditions^76,77^, supporting that generalist or stress-tolerant species may also act as connectors, facilitating interactions in challenging environments^75,77^. Furthermore, AMF presented relatively few connections to oil palm nodes, suggesting that AMF is more specialized within the network. This supports growing evidence that intensive land-use practices in oil palm monocultures may negatively impacts AMF diversity^24,78^.

Global environmental changes inevitably interfere with biotic interactions, driving the successful establishment and dispersion of non-native taxonomic groups^35,37,38,79,80^. Key processes or drivers of successful invasions include disturbance due to physical or chemical resource flux, such as climate and fertilization^37,38^. These external drivers act as environmental filters, favoring adaptable and opportunistic species that occupy a broader range of ecological niches, ultimately outcompeting the resident species^38^. Non-native species have specific characteristics that promote resilience and may favor successful seedling recruitment^79,80^. Traits such as long-distance dispersal, reproductive plasticity, ability for nitrogen fixation, and allelopathy are competitive advantages over native species that promote seed persistence even in unfavorable conditions^35,37,38,79,80^. As a result, this interference gives them a competitive edge over native species that rely on mutualistic relationships. For example, the invasive perennial shrub *C. hirta,* identified in this study as an indicator species of oil palm monocultures, has a rapid growth rate, prolific seed production, shade tolerance, and it releases allelochemicals into the soil that gives it a competitive advantage over the native flora^79^. Besides that, this species is tolerant to extreme environmental conditions, inhabiting disturbed areas, and thriving in modified open habitats such as roadsides and tree plantations^4,66,79,80^. Furthermore, the non-native climbing herb *A. gangetica* has been reported to exert a detrimental impact on oil palm plantation yields. It demonstrates rapid and expansive dispersion and possesses a generalist nature that allows it to adapt to various types of soil. Both herbs compete with crops and native species for resources^81,82^. Furthermore, the presence of opportunistic non-native plant species may influence the diversity of the soil microbiota by promoting soil changes through enzymatic processes and direct or indirect effects on carbon and nitrogen cycles^38^.

### Land-use effects on network connectivity

Indicator species of healthy or less-converted systems have shown higher connectivity within the network^8^, underscoring the importance of assessing root-associated microbiota. In addition, indicator taxa associated with the agroecosystem jungle rubber, which plays a role as a transitional zone between mildly disturbed and disturbed habitats, encompasses indicator taxa associated with both types of habitats. For instance, Glomeraceae spp. and Herpotrichiellaceae spp. were identified as indicators of jungle rubber, with both AMF and stress-resistant groups playing significant roles in shaping the network^21,27,83^. Plant and fungal species identified as indicator species of rainforests likely exert a substantial influence on resource flow. Consequently, the removal of these species can result in significant alterations to community composition and ecosystem functioning^46^. In this study, the tree indicator species identified from rainforests belonged to the orders Ericales, Gentianales, Laurales, Magnoliales, Malpighiales, Myrtales, Santalales, and Sapindales. Identifying a wide range of indicator species associated with specific land-use types provides valuable insights for restoration efforts and could be further explored through biodiversity enrichment experiments^84^.

Conservation management and restoration of ecosystem functions and services may be achieved by focusing on interaction networks and targeting indicator species in order to leverage ecosystem stability^85^.

## Conclusion

Our study is a pioneering effort to explore plant roots and their connection with the fungal community in the framework of forest conversion systems using plant and fungal barcodes from identical DNA samples. Based on our analysis, we have observed coexistence between the cultivated crop plants and generalist or stress-resistant organisms. Interestingly, agroforestry systems may function as an intermediate zone between mildly disturbed and highly disturbed habitats, hosting indicator fungal species characteristic of both habitat types.

We identified topological and structural differences among the fungal and plant communities in different land-use types. Modules connected to less managed land-use types displayed enhanced connectivity and harbored indicator species associated with rainforest or jungle rubber systems. This contrasted with modules containing indicator species linked to plantation plots. Moreover, rare or endemic taxa frequently played a crucial part in the network structure by forming multiple connections between modules that shared indicator species associated with rainforest habitats. This highlights the impact of monocultures on the composition and structure of plant-fungal networks and points out the potential role of agroforestry systems in upholding diversity, connections, and, subsequently, ecosystem functioning.

